# MitoChontrol: Adaptive mitochondrial filtering for robust single-cell RNA sequencing quality control

**DOI:** 10.64898/2026.04.04.716517

**Authors:** Caitlin Strassburg, Danielle Pitlor, Aatur D. Singhi, Rachel A. Gottschalk, Shikhar Uttam

## Abstract

**Summary:** Mitochondrial transcript abundance is a standard quality control metric in single-cell RNA sequencing, but fixed percentage thresholds fail to account for the substantial variation in mitochondrial content across cell types and tissues, risking both retention of compromised cells and exclusion of transcriptionally active viable cell populations. We present MitoChontrol, a cell-type-aware probabilistic framework for mitochondrial quality control that models the mitochondrial transcript fraction within transcriptionally coherent clusters as a Gaussian mixture distribution. Compromised-cell components are identified from the upper tail of each cluster-specific distribution, and filtering thresholds are defined as the point at which the posterior probability of cellular compromise exceeds a user-definded confidence value. Applied to controlled perturbation experiments and a pancreatic ductal adenocarcinoma single-cell dataset, MitoChontrol selectively removes transcriptionally compromised cells while preserving biologically elevated but viable populations, outperforming fixed-threshold and outlier-based approaches.

**Availability and Implementation:** MitoChontrol is implemented in Python and integrates directly with AnnData-based workflows. It is freely available under the GNU General Public License v3 (GPL-3.0) at: https://github.com/uttamLab/MitoChontrol (DOI: https://doi.org/10.5281/zenodo.19423054)

## 1. Introduction

Single-cell RNA sequencing (scRNA-seq) is a high-throughput technology for characterizing the transcriptomes of individual cells at scale. It has enabled high-resolution dissection of cellular heterogeneity, revealing distinct cell types, transient states, cell fate decisions, and intercellular interactions in the context of development, tissue homeostasis, and disease Tanay and Regev (2017); Antkowiak et al. (2025). However, scRNA-seq data are susceptible to technical artifacts that can confound downstream analyses and biological interpretation Luecken and Theis (2019); Kim et al. (2015).

A common source of technical artifact is cellular compromise arising during sample preparation Slyper et al. (2020); Stegle et al. (2015). Disruption of membrane integrity leads to cytoplasmic RNA leakage and the induction of cellular stress responses, collectively resulting in an elevated relative abundance of mitochondrial transcripts (mtRNA) Galluzzi et al. (2012). Accordingly, the mitochondrial transcript fraction is a widely used quality control (QC) metric, and cells exceeding a defined threshold are typically excluded prior to downstream analysis Ilicic et al. (2016).

The conventional approach applies a fixed threshold — most commonly 10% — to the mitochondrial transcript fraction Osorio and Cai (2020). However, mitochondrial content can vary substantially with metabolic demand in a cell-type- and tissue-specific manner Subramanian et al. (2022); Ying et al. (2026). Fixed thresholds are therefore poorly suited to balancing the competing objectives of removing compromised cells and preserving biologically meaningful heterogeneity. A threshold set too permissively retains compromised cells; one set too stringently excludes metabolically active but viable cells, and in datasets with pronounced metabolic diversity, may eliminate entire cell populations. This is particularly consequential in tumor microenvironments, where malignant and stromal cell populations frequently exhibit significantly elevated mitochondrial activity relative to other cell types Yates et al. (2025); Du et al. (2025). Fundamentally, fixed thresholding assumes a homogeneous mitochondrial distribution, and therefore is unable to account for biological variability across cell populations.

Adaptive methods have been developed to address this limitation. miQC Hippen et al. (2021) models the joint distribution of mitochondrial transcript fraction and the number of detected genes — another standard QC metric — using a two-component mixture of linear regression models, one representing intact cells and one representing compromised cells, to assign each cell a posterior probability of compromise. This probabilistic framework allows thresholds to adapt to the dataset-wide bimodal distribution arising from the co-occurrence of healthy and compromised cells. However, miQC fits a single global model in which the intact-cell component implicitly assumes that healthy cells form a coherent distribution in the gene-count/mtRNA space. In datasets containing cell types with meaningfully different baseline mitochondrial activity, this assumption does not hold: the healthy population is itself a mixture of cell-type-specific distributions, and cells from high-mtRNA cell types can be drawn toward the compromised component not because they are damaged but because their baseline mitochondrial content exceeds the global expectation. miQC therefore has limited capacity to resolve cell-type-specific metabolic heterogeneity. ddQC Subramanian et al. (2022) takes a complementary approach by first grouping cells into clusters of similar transcriptional profiles and then applying cluster-specific, median absolute deviation (MAD)-based thresholds, allowing QC criteria to reflect cell-type-associated differences in mitochondrial content. However, as an outlier detection approach, ddQC assumes an approximately unimodal mtRNA distribution within each cluster, and is not robust to distributional diversity that may arise due to the co-ocurrence of compromised and healthy cell populations.

To address these concerns, we introduce MitoChontrol, a cell-type-aware mitochondrial filtering method that models mitochondrial transcript fractions using mixture distributions to explicitly represent distinct transcriptional states, enabling identification of compromised cells and inference of interpretable, dataset-specific thresholds based on the posterior probability of cellular compromise.

## 2. MitoChontrol

### 2.1. Model

MitoChontrol models the distribution of mitochondrial transcript fractions within transcriptionally coherent cell populations to infer dataset-specific filtering threshold based on the posterior probability level of compromise. Cells are first stratified into clusters of transcriptionally similar cells (see Section 2.2). Within each cluster, the mitochondrial RNA fraction is modeled as a Gaussian mixture distribution,

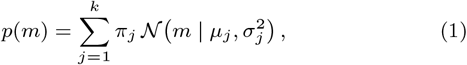

where *m* denotes the mitochondrial transcript fraction, *k* is the number of mixture components, and *π*_*j*_, *µ*_*j*_, and 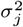 denote the weight, mean, and variance of component *j*, respectively. Models are fit for *k* = 1, …, *k*_max_ (default *k*_max_ = 5), and the optimal number of components is selected using the Bayesian Information Criterion (BIC) by balancing model complexity while capturing intra-cluster variation. Parameter estimation is performed using an online expectation–maximisation (EM) algorithm to enable efficient fitting on large single-cell datasets.

The filtering threshold is computed probabilistically rather than by a fixed mtRNA percentage cutoff. First, components representing cellular compromise are identified by examining the upper tail of the empirical distribution – corresponding to highly elevated mitochondrial content – where compromised cells are expected to accumulate. Components contributing to distribution tail even with a probability of 1% are designated as compromised-cell components. Next, the posterior probability of compromise at a given mtRNA value *m* is computed as the summed posterior probability of membership across these components,

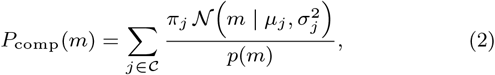

where *m* denotes the set of compromised components and *p*(*m*) is defined in Equation 1. A filtering threshold *m*^*^ is defined as the lowest mtRNA fraction value at which *P*_comp_(*m*) exceeds a user defined confidence level (default *τ* = 0.8),

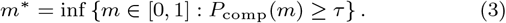

Cells whose mitochondrial transcript fraction exceeds *m*^*^ are classified as compromised and excluded from downstream analysis.

If no mtRNA value satisfies the confidence criterion – indicating that no distinct compromised population is identifiable within the cluster – the user can select a fallback strategy from three options: apply the conventional threshold of 10% mtRNA, keep entire cluster, or filter out the entire cluster.

This probabilistic formulation allows mitochondrial filtering thresholds to adapt to the cluster-specific distributional structure of mtRNA, while preserving biologically meaningful variation in mitochondrial activity across cell types.

### 2.2. Implementation

MitoChontrol is implemented as a Python package designed to integrate directly with standard single-cell RNA-seq analysis workflows.

#### Cell-type stratification

To condition mitochondrial filtering on transcriptional context, cells are first partitioned into transcriptionally homogeneous clusters. Raw count matrices are processed using a standard single-cell preprocessing workflow in R including ambient RNA removal Lun et al. (2019), doublet detection McGinnis et al. (2019), gene filtering, and normalization Hao et al. (2021). Count matrices are converted to *H5AD* format Marsh et al. (2025) before being loading as an AnnData object in Python. Dimensionality reduction is performed using principal component analysis with *p* (default *p* = 30) components, and clusters are identified using Leiden community detection on a *k*-nearest neighbor graph Wolf et al. (2018). Clustering resolution is automatically selected using resolution tuning that balances silhouette score and modularity to maximize separation while avoiding excessive fragmentation. Clusters are treated as independent strata for downstream modeling.

#### Mixture modeling

For each cluster, the fraction of mitochondrial transcripts per cell are extracted from raw counts and modeled using the Gaussian mixture framework described above (See 2.1). Model parameters are initialized using *k*-means, quantile-based initialization, or random sampling. Mixture parameters are then optimized using an online expectation–maximization algorithm with a fixed number of passes through the data (default *T* = 3).

#### Posterior probability thresholding

Posterior probabilities of component membership are evaluated across the range of observed mtRNA values. Compromised components associated with elevated mtRNA are automatically identified by examining posterior mass in the upper tail of the distribution. Cells are classified as compromised when the posterior probability of belonging to these components exceeds a user-defined confidence level. A high default confidence level of 0.80 probability of compromise is recommended, but may be decreased when compromised components are easily separable.

#### Outputs and integration

MitoChontrol takes an AnnData object as its input, and outputs cluster-specific mtRNA thresholds and cell-level filtering labels, which are appended directly to the input AnnData object. These annotations are preserved when exported to *H5AD* format, enabling interoperability with a broad ecosystem of single-cell analysis tools across programming environments. Optional diagnostic outputs include mixture model fits, posterior probability curves, and cluster-specific threshold visualizations to facilitate quality assessment. All outputs are designed to integrate seamlessly with existing single-cell analysis workflows without requiring additional data restructuring.

#### Time complexity

Mixture modeling is performed independently for each cluster. For a cluster containing *n* cells, fitting a *k*-component model via online EM requires *O* (*nTk*) operations, where *T* is the number of EM passes. Because models are evaluated for *k* = 1, …, *k*_max_, the total cost per cluster is 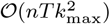. For fixed *k*_max_ and *T*, runtime scales approximately linearly with the number of cells.

## 3. Results

### MitoChontrol infers adaptive mtRNA thresholds from mixture models

Figure 1 illustrates how MitoChontrol derives data-driven mitochondrial RNA (mtRNA) filtering thresholds using a probabilistic model rather than a fixed mitochondrial percentage cutoff. For each cell cluster, mtRNA fractions are modeled using a Gaussian mixture with number of components selected via Bayesian Information Criterion (BIC) (see 2.2). Figure 1A illustrates a representative cluster in which the empirical mtRNA distribution exhibits a pronounced right-tailed structure. The selected mixture model captures this structure with distinct components corresponding to baseline and elevated mtRNA levels.

**Figure 1:**
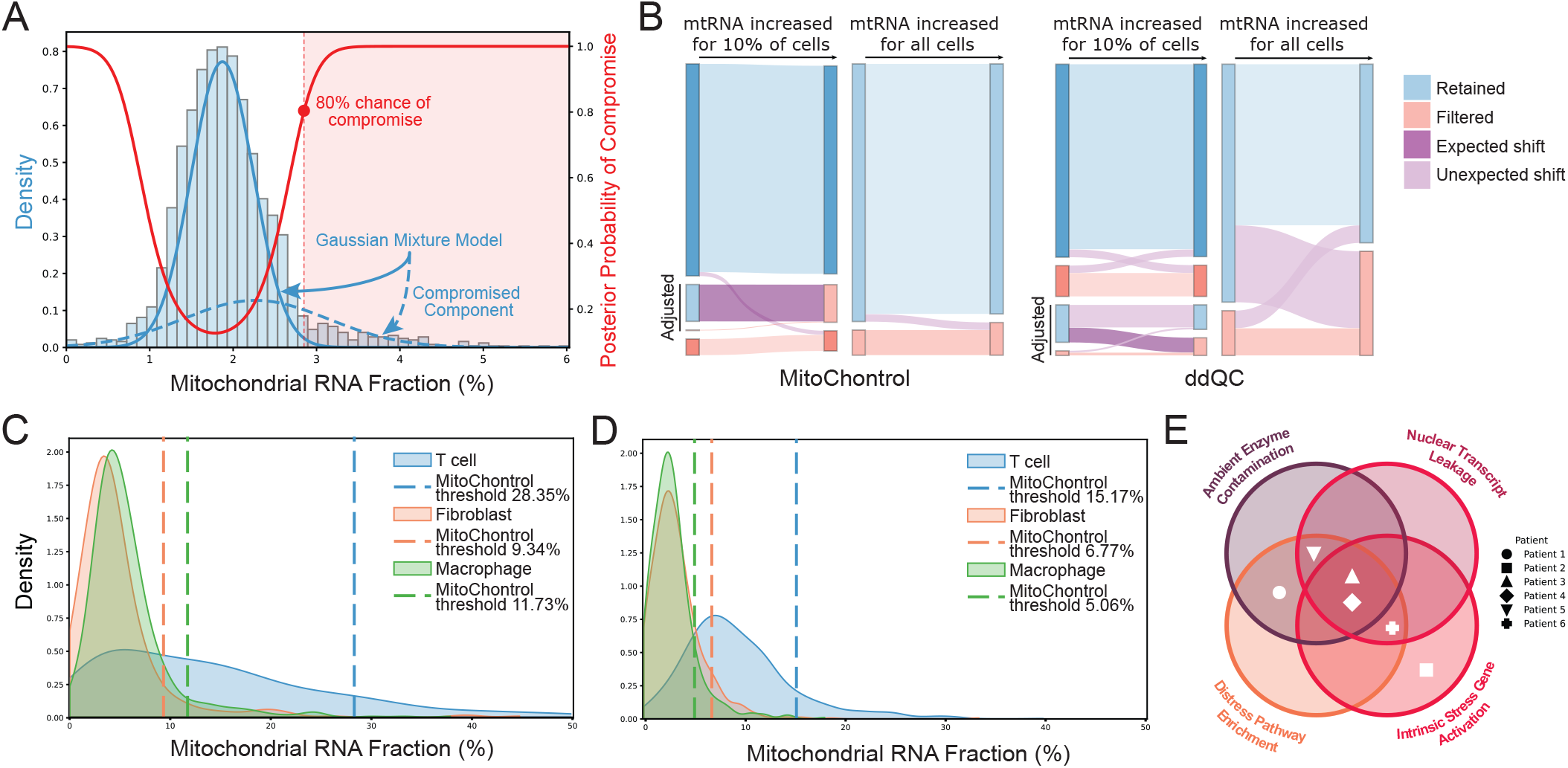
MitoChontrol adaptively identifies compromised cells using probabilistic modeling of mitochondrial transcript fractions. (**A**) Example Gaussian mixture model fit to mitochondrial RNA (mtRNA) fractions within a transcriptionally coherent cluster. The empirical mtRNA distribution (histogram) is modeled using a BIC-selected Gaussian mixture (blue). Posterior probabilities of component membership are evaluated across the mtRNA range, and the filtering threshold is defined as the lowest value at which the posterior probability of compromise (red) exceeds a predefined confidence cutoff (80(**B**) Benchmarking under simulated perturbations. A baseline distribution of healthy HEK293 cells was subjected to two controlled perturbations: localized compromise (mtRNA increased in 10% of cells) and global metabolic shifts (mtRNA increased uniformly across all cells). Expected changes in classification following perturbation are highlighted in dark purple, erroneous changes in light purple. MitoChontrol selectively removes the compromised subset while preserving the baseline distribution, whereas ddQC exhibits reduced specificity under these conditions. (**C–D**) Application to pancreatic ductal adenocarcinoma (PDAC) single-cell RNA-seq data. Cluster-specific mtRNA distributions and inferred thresholds are shown for representative immune and stromal populations in tumor tissue (C) and adjacent noncancerous tissue (D). Thresholds vary substantially across cell types, illustrating the limitations of a single global mtRNA cutoff. (**E**) Evidence of cellular compromise in filtered cells. Filtered populations were evaluated for transcriptional signatures associated with cellular distress, including ambient enzyme contamination, nuclear transcript leakage, intrinsic stress responses, and enrichment of distress-associated pathways. Across patients, filtered cells consistently exhibited signatures corresponding to at least one distress axis.

Components associated with elevated mtRNA are automatically selected as compromised-cell components and the posterior probability of compromise is evaluated across the mtRNA range (see 2.1). Cells with a probability of compromise exceeding a predefined confidence level cutoff (default *τ* = 0.80) are filtered from the sample. This procedure yields cluster-specific mtRNA thresholds determined by the shape of the observed distribution. Importantly, applying the same posterior probability cutoff across clusters produces different mtRNA percentage thresholds, reflecting cluster-specific tail structure opposed to a uniform cutoff.

### MitoChontrol distinguishes compromised cells from cell state associated shifts

To establish the efficacy of MitoChontrol, we compared its performance with that of ddQC Subramanian et al. (2022) by applying both methods to specific and controlled synthetic perturbations of mtRNA content in an experimental scRNA-seq dataset, designed to assess the effectiveness and sensitivity of the methods in filtering out compromised cell proportions (Fig. 1B). The adaptive QC method miQC Hippen et al. (2021) was also evaluated; however, the simulated perturbations violated the joint mtRNA–gene count assumptions required by its model, preventing stable threshold estimation. Specifically, an scRNA-seq dataset of healthy FACS-sorted HEK293 cells served as the baseline distribution Ordoñez-Rueda et al. (2020). Data was accessed from ENA repository: https://www.ebi.ac.uk/ena with the study accession number PRJEB33078.

To simulate cellular compromise, mtRNA fractions were increased in 10% of the cells, producing a right-tailed expansion of the mtRNA distribution (Fig. 1B, left). Under this perturbation, MitoChontrol preferentially removed the inflated subset while overwhelmingly preserving the baseline population. In contrast, ddQC retained many cells from this subset while also excluding some unperturbed cells.

When the mtRNA perturbation was extended to 100% of the cells (Fig. 1B, right), simulating increased mitochondrial activity rather than compromise, MitoChontrol adjusted the inferred threshold accordingly, resulting in minimal change to the retained and filtered populations. ddQC did not fully adjust to the shifted distribution, leading to increased filtering of otherwise unchanged cells.

Taken together, these perturbations illustrate how MitoChontrol is able to identify increase in compromised cell population and filter them, while also being able to distinguish it from overall shifts in mitochondrial transcript abundance.

### MitoChontrol adaptively filters compromised cells in pancreatic cancer

To evaluate MitoChontrol in a biologically heterogeneous context, we applied the method to a pancreatic ductal adenocarcinoma (PDAC) scRNA-seq dataset Chen et al. (2023). Data was accessed from GEO Repository with Accession Number: GSE212966 (PRJNA879876). We began by preprocessing and clustering the data as described in 2.2, yielding cell-type clusters representing diverse immune and stromal populations including B cells, T cells, NK cells, plasma cells, monocytes, neutrophils, macrophages, dendritic cells, fibroblasts, endothelial cells, epithelial cells, and endocrine cells spanning many unique activation states.

For each of these cell-type clusters, MitoChontrol identified distinct confidence-defined thresholds. This is demonstrated via representative T, fibroblast, and macrophage cell populations that illustrate MitoChontrol’s adaptive approach across stromal and immune (both innate and adaptive) cells in PDAC tumor (Fig. 1C) and adjacent noncancerous tissue (Fig. 1D); MitoChontrol inferred thresholds adapted to each cell type’s mtRNA distribution (Table 1).

**Table 1.** MitoChontrol adaptively selects cell-type specific mtRNA thresholds in pancreatic ductal adenocarcinoma scRNA-seq data.

| Cell Type | Cancerous Tissue | Adjacent Tissue |
| --- | --- | --- |
| T cell | 28.53% | 15.17% |
| Fibroblast | 9.34% | 6.77% |
| Macrophage | 11.73% | 5.06% |

Critically, MitoChontrol-filtered cells exhibited transcriptional signatures consistent with cellular compromise, confirmed by differential expression and pathway enrichment analyses Fang et al. (2023) of filtered and retained cells within clusters.^1^ Among nine samples with significant differential expression (*p*_val_ < 0.05, log2 fold-change > 2), the filtered fraction of seven samples showed enrichment of *MALAT1*, a transcript previously associated with low-quality scRNA-seq libraries Clarke and Bader (2024). Filtered cell populations across the cell types were also pooled across all patients, and reanalyzed by classifying them as exhibiting compromise when they showed enrichment of at least one distress axis, including ambient enzyme contamination, nuclear transcript leakage, intrinsic cellular stress responses, or enrichment of distress-associated pathways. Across patients, filtered cells consistently exhibited signatures corresponding to one or more of these categories.(Fig. 1E).

We note that since MitoChontrol thresholds are cell type specific, cell populations with naturally elevated mitochondrial fractions were not systematically penalized, and only those cell populations exhibiting pronounced right-tailed structure underwent more stringent filtering. Together, these results demonstrate that MitoChontrol adaptively filters cells with transcriptional signatures consistent with compromise while preserving transcriptionally active but not-compromised cell populations across heterogeneous tumor microenvironments.

## 4. Discussion

Mitochondrial transcript abundance is widely used as a QC metric in single-cell RNA sequencing, yet the common practice of applying fixed mtRNA thresholds implicitly assumes homogeneous transcription of mtRNA across cell populations. This assumption is frequently violated in heterogeneous tissue microenvironments, where mitochondrial content varies with cell type, activation state, and microenvironment Subramanian et al. (2022); Du et al. (2025). By first identifying transcriptionally coherent cell populations, and then probabilistically modeling mtRNA distributions within each population as mixture models, MitoChontrol reframes mitochondrial filtering as a probabilistic inference problem, providing a more nuanced threshold control. Furthermore, since MitoChontrol models the mtRNA fractions independently of other QC metrics, it can be integrated with complementary QC tools. This modularity is particularly valuable as single-cell preprocessing pipelines continue to evolve.

We have demonstrated that MitoChontrol provides an adaptive way to address the challenge of identifying and filtering compromised cells, but quality control of scRNA-seq remains a complex task Stegle et al. (2015). In clusters with very small cell counts or minimal distributional separation, mixture components may be difficult to resolve, and conventional heuristics may need to be used. Additionally, mitochondrial transcript abundance reflects multiple biological processes beyond cellular compromise, including metabolic activation and mitochondrial biogenesis Galluzzi et al. (2012). Although, as shown, MitoChontrol is able to avoid unnecessary filtering in cases of global shift, users should always examine the filtered cells for confirmation.

## 5. Conclusion

MitoChontrol formalizes mitochondrial RNA filtering as a cluster-specific probabilistic inference problem, and helps ameliorate limitations of fixed-percentage thresholding which fail to capture the biological complexity of mtRNA expression. By modeling mtRNA distributions with BIC-selected mixture models and defining thresholds through posterior confidence, the method adaptively identifies compromised cells. In both controlled perturbation experiment and the heterogeneous scRNA-seq PDAC data, MitoChontrol selectively removes transcriptionally compromised cells while reducing cell-type retention bias. This distribution-aware framework provides a statistically principled and computationally lightweight alternative to conventional mtRNA cutoff-based quality control in single-cell RNA sequencing workflows.

## 6. Acknowledgments

The authors thank the reviewers for their valuable suggestions.

## 7. Author contributions

Caitlin Strassburg (Conceptualization [equal], Formal analysis [equal], Methodology [equal], Resources [equal], Software [lead], Writing—review & editing [equal]); Danielle Pitlor (Resources [equal]); Aatur D. Singhi (Resources [equal]); Rachel A. Gottschalk (Conceptualization [equal], Methodology [equal], Resources [supporting], Funding acquisition [equal], Writing—review & editing [equal], Project supervision [equal]); Shikhar Uttam (Conceptualization [equal], Formal analysis [equal], Methodology [equal], Resources [lead], Software [Supporting], Software-review [lead], Funding acquisition [equal], Writing—review & editing [equal], Project supervision [equal])

## 10. Data Availability

No new data were generated or analysed in support of this research.

## Footnotes

1 We note that interpretation of compromised-cell expression profiles is inherently limited by small sample sizes.

